# OPM-MEG reveals protracted maturation of visual phase-amplitude coupling from early childhood to adulthood

**DOI:** 10.64898/2026.09.08.749145

**Authors:** Lyam M Bailey, Natalie Rhodes, Robert A Seymour, Matthew J Brookes, Margot J Taylor

## Abstract

Cross-frequency coupling describes interactions between neural ensembles operating at different frequency bands, and has been proposed to underlie coordination of brain activity over multiple temporal and spatial scales. Phase amplitude coupling (PAC) describes modulation of the amplitude of high-frequency oscillations by the phase of low frequency oscillations. MEG work in adults has demonstrated that alpha-gamma PAC may support visual perception. However, very little work has investigated visual PAC in children, or the development of this process between childhood and adulthood. Here we investigated the development of PAC in a sample of 101 individuals, comprising 50 children (aged 2-13 years) and 51 adults (21-34 years). Participants completed a passive visual stimulation paradigm while neural responses were recorded using a wearable whole-head OPM-MEG system. Sensor-level data were spatially resolved to visual cortex, and we quantified coupling between the phase of low-frequency oscillations (5-13 Hz) and the amplitude of broadband gamma oscillations (28-100 Hz) in each participant. In the adult data, we identified a cluster of phase-amplitude frequency pairs (phase: 7-11 Hz, amplitude: 52-66 Hz) in which there was strong evidence for an increase in PAC following stimulus presentation relative to baseline. Within this cluster, the magnitude of stimulus-induced PAC was positively correlated with age across all participants. Meanwhile, we did not see credible evidence for visual PAC at any frequency pair(s) within the child group. These results suggest that, alongside well-documented developmental changes in band-specific oscillatory power and connectivity, visual PAC represents a distinct functional process which matures with age.

## 1. OPM-MEG reveals protracted maturation of visual phase-amplitude coupling from early childhood to adulthood

Non-invasive neurophysiological imaging techniques such as magneto- and electroencephalography (M/EEG) are invaluable tools for understanding functional maturation of the developing brain. In particular, owing to their millisecond-level temporal resolution, these techniques are useful for assessing neural oscillations underlying perception and cognition. Much developmental MEG research has focused on age-related changes in long-range connectivity and/or local (region-specific) oscillatory power within physiologically-relevant frequency bands (see Edgar et al., 2026; Rhodes et al., 2024 for discussions). In addition, recent years have seen growing interest in a complementary form of oscillatory-based neural activity, known as cross-frequency coupling. Cross-frequency coupling refers to interactions between neural ensembles operating at different frequency bands (Canolty & Knight, 2010); as such, it can capture local connectivity supporting hierarchical information processing (e.g., between populations at different cortical layers; Bonaiuto et al., 2018; Sotero et al., 2015; and/or spatial scales; Canolty & Knight, 2010).

One form of cross-frequency coupling is phase amplitude coupling (PAC), whereby the amplitude of oscillations at high frequencies (typically in the beta band: 13-30 Hz, or gamma band: 30-100 Hz) is modulated by the phase of oscillations at lower frequencies (theta: 4-7 Hz: alpha: 7-13 Hz). Work in typical adult populations has implicated PAC in a large range of perceptual and cognitive domains: from low-level visual (Seymour et al., 2017; Voytek et al., 2010), auditory (Li et al., 2025), and sensorimotor processing (Das et al., 2025; Li et al., 2026), to working memory (Daume et al., 2024) and spoken language comprehension (Lizarazu et al., 2023; Oderbolz et al., 2024; Weissbart & Martin, 2024). Given the apparent ubiquity of cognitively-relevant PAC in the adult brain, efforts to characterize its developmental trajectories may yield important advances in our broader understanding of functional brain maturation.

A number of studies have investigated PAC effects in children and infants (e.g., Barnes et al., 2016; Kang et al., 2026; Kroupi et al., 2024; Mamashli et al., 2018; Mariscal et al., 2021; Peck et al., 2022; Port et al., 2019; Tokariev et al., 2016, 2022), with age-related increases seen from as young as three months of age (Mariscal et al., 2021). However, these studies have mainly depended on resting-state or sleep data, while task-related PAC studies in children are (to our knowledge) sparse. We note that Mamashli et al. (2018) investigated the development of alpha-gamma PAC in the fusiform gyrus, elicited by images of faces, and reported a significant increase between childhood (ages 7-13) and adolescence, while Barnes et al. (2016) reported that longitudinal cognitive training resulted in increased in cross-region (frontal-temporal and parietal-temporal) alpha-gamma PAC in the context of a spatial working memory task, in children aged 8-11. These studies indicate that cognitively relevant PAC is at least present in school-age children and may increase with age and/or cognitive ability.

Here we investigated the developmental trajectory of PAC supporting visual processing in a cohort of children and adults, aged 2-34 years, using optically pumped magnetometers (OPMs). We expected that visual PAC would exhibit age-related changes between childhood and adulthood as there is a great deal of evidence that responses to visual stimuli in the alpha and gamma frequency bands (the constituent bands of alpha-gamma PAC observed in adults; Seymour et al., 2017; Voytek et al., 2010) undergo significant maturational changes over the same developmental period. In particular, peak frequency in the alpha band and power in the gamma band both increase with age, while the frequency distribution of the gamma response differs between children and adults (Cellier et al., 2021; Gaetz et al., 2011; Miskovic et al., 2015; Orekhova et al., 2018; Rhodes, Rier, Singh, et al., 2025; Vandewouw et al., 2024). As such, we expected that visual PAC would similarly mature with age; manifesting either as a shift in frequency ranges at which PAC appears between childhood and adulthood, or as increasing PAC magnitude within the same frequency ranges. As an aside, we note that visual PAC has been observed in a group of mid-to-late adolescents and young adults (aged 14-20; Seymour et al., 2019); whether visual PAC is present at younger ages remains unclear.

## 2. Methods

### 2.1. Participants

We analyzed an OPM-MEG dataset previously described in Rhodes, Rier, Singh, et al. (2025). In brief, this dataset was acquired from 51 children (2-13 years of age) and 51 adults (21-34 years) who underwent scanning at The Hospital for Sick Children, Toronto, Canada (SK; 24 children, 26 adults), or at the Sir Peter Mansfield Imaging Centre, University of Nottingham, UK (UoN; 27 children, 26 adults). One child in the UoN cohort was excluded from analyses due to incomplete 3D head digitization while wearing the OPM helmet (required for coregistration; see Data Acquisition), therefore we analyzed data from 50 children and 51 adults. All participants or their parents or legal guardians provided written informed consent; all children provided verbal assent. All procedures were approved by a local institutional research ethics board at each site.

### 2.2. Paradigm

Participants completed a passive visual stimulation paradigm. Each trial began with a central fixation cross presented for 1250 ms (+/- 200 ms jitter) followed by an inwardly moving circular grating for 1000 ms. The circular gratings moved at 1.2°s^-1^ with 1.32 cycles per degree and subtended a visual angle of 7.6°. Each participant underwent 60 visual stimulation trials, interspersed with images of human faces and cartoon characters (not analysed).

### 2.3. Data Acquisition

Data were collected from two international OPM-MEG systems, situated at SK and UoN. Technical specifications regarding each system, as well as the magnetically-shielded rooms (MSRs) in which they were housed, are provided in Rhodes, Rier, and Singh et al. (2025). Previous work has demonstrated equivalent performance between these two systems (Hill et al., 2022), while Rhodes, Rier, and Singh et al. (2025) reported no significant between-site differences in visual gamma responses within the age- and sex-matched adult subsample of the dataset analyzed by the present study. These findings suggest that between-site differences were unlikely to confound our analyses.

Briefly, the SK system comprised an 80-channel array (from 40 dual-axis OPMs, 3rd- generation QZFM; QuSpin, Colorado, USA) and the UoN system consisted of a 192- channel array (64 triaxial OPMs, 3rd-generation QZFM). Both sensor arrays were mounted in rigid 3D-printed helmets (Cerca Magnetics Ltd. Nottingham, UK), and their spatial layouts were similarly configured to provide good coverage of visual cortex. Sensors were mounted in one of five helmet sizes to accommodate a range of head sizes across the age range. Participants were seated at the centre of the MSR between bi-planar coils which provided active magnetic field control (Holmes et al., 2019). OPM data were recorded at 1200 Hz using a National Instruments (NI, Texas, US) data acquisition system interfaced with LabView (NI). Data were recorded by a Windows PC connected to the OPM system; timings of experimental events (i.e., stimulus onsets / offsets) were sent to the recording PC via a parallel port. At both sites, visual stimuli were projected onto a non-magnetic screen via an MEG-compatible visual stimulus back projection system; participants were seated at a viewing distance of approximately 1 m from this screen.

An optical imaging system (EinScan H, SHINING 3D, Hangzhou, China) was used to obtain two 3D head digitizations from each participant (one with the OPM helmet; one without). These 3D images were used to generate a ’pseudo-MRI’ for each participant (obtained by warping an age-matched template MRI to their digitized head image; Rhodes, Rier, Boto, et al., 2025), which in turn was used for coregistration of sensor positions and orientations to brain anatomy. Head digitations without the helmet (required for warping) could not be acquired for 20 children; in these cases, sensor data were coregistered directly to age-matched template MRIs.

### 2.4. Data Preprocessing and Beamforming

We used preprocessed and spatially resolved data from Rhodes et al. (see Rhodes, Rier, Singh, et al., 2025 for a full description of preprocessing and beamforming procedures; relevant code can be found at https://github.com/nsrhodes/gamma_opm_2024). Briefly, preprocessing included bad channel removal, notch filtering at power line frequencies and their harmonics (UoN: 50 Hz, SK: 60 Hz), band-pass filtering (1-150 Hz), homogenous field correction (Tierney et al., 2021), and epoching. Epochs were 3 s in length and included 1 s before and 2 s following the onset of visual gratings. Additional preprocessing included manual trial rejection and ICA to remove eye blinks and cardiac artefacts. Preprocessed, epoched data were beamformed to a virtual electrode at the location of the peak visual gamma response for each subject. This location was determined by contrasting gamma power (30-80 Hz range) following stimulus presentation (0.3 to 1 s relative to stimulus onset) to that of the preceding fixation period (-0.8 to –0.1 s). This contrast provided a pseudo-T statistical image; this image was constrained to a visual cortical mask, and the position of the peak T value within the mask was used as the position of the virtual electrode.

Lead fields were computed from a single-shell forward model (Nolte, 2003), fitted to each participant’s pseudo-MRI image. We conducted our analyses on these functionally-defined virtual electrodes. We note that the spatial position of peak gamma responses did not vary systematically with age (Rhodes, Rier, Singh, et al., 2025); therefore our age-related analyses were unlikely to be affected by developmental changes in peak location.

### 2.5. Phase-Amplitude Coupling (PAC) Analysis

We investigated stimulus-induced phase-amplitude coupling (PAC) in the epoched, spatially resolved data based on the procedure described in Seymour et al. (2017). PAC analyses were conducted in the MATLAB environment using custom scripts adapted from publicly available code (https://github.com/neurofractal/sensory_PAC).

We investigated coupling between phase in the alpha band (defined as 5-13 Hz) and amplitude in the gamma band (28-100 Hz). Alpha is typically defined as 7-13 Hz; we extended the lower bound of this range by 2 Hz to account for the fact that peak alpha frequency can be lower than 7 Hz in young children (e.g., Cellier et al., 2021; Vandewouw et al., 2024), potentially leading to PAC effects at low phase frequencies. In addition, Rhodes, Rier, and Singh et al. (2025) found that young children exhibit a peak visual gamma response around 30 Hz. We reasoned that age-related PAC effects may also occur at this frequency, motivating us to extend the gamma band range to 28 Hz.

We estimated PAC in each cell of a *f*_p_ × *f*_a_ grid comprising low (alpha) frequencies carrying phase information, *f*_p_, and high (gamma) frequencies carrying amplitude information, *f*_a_. To obtain *f*_p_ and *f*_a_ we first filtered the data to their corresponding frequencies using fourth-order, two-pass Butterworth filters^1^. Filter bandwidth for extracting *f*_p_ was fixed at the centre frequency ± 1 Hz, while the bandwidth for extracting *f*_a_ was defined as the centre frequency ± (centre frequency × 0.4). *f*_p_ and *f*_a_ were extracted in 1 Hz and 2 Hz steps respectively. After filtering, phase (angle) and amplitude components were extracted from *f*_p_ and *f*_a_ respectively using Hilbert transforms; these were used to compute our PAC metrics (see below). It is worth noting that robust PAC estimation requires the *f*_a_ bandwidth to be wide enough to capture modulating *f*_p_ oscillations, i.e., the filter sidebands must be ≥ *f*_a_ ± *f*_p_ (Aru et al., 2015). As our *f*_a_ filter sidebands were defined as the centre frequency × 0.4, PAC estimates at the lower end of our gamma range (< 34 Hz) would not be meaningful at higher *f*_p_ frequencies. We therefore excluded computed PAC estimates at the intersection of *f*_p_ [10, 13 Hz] and *f*_a_ [28, 34 Hz] from statistical analyses and figures.

We quantified coupling between phase and amplitude in each *f*_p_, *f*_a_ pair using the phase-locking-value modulation index (PLV-MI) described in Cohen et al. (2008). The PLV-MI assumes that, if PAC is present, the *f*_a_ amplitude envelope will oscillate at a modulating *f*_p_ frequency. Hilbert transforms are used to obtain the instantaneous angle (i.e., phase) of *f*_p_ and amplitude envelope of *f*_a_. Next, the phase of the (Hilbert-derived) *f_a_* amplitude envelope is extracted by a second Hilbert transform. The PLV-MI is quantified by a phase locking value which describes the magnitude of phase synchronization between the phase of the *f_a_* amplitude envelope and the phase of *f*_p_. To ensure that results were not biased by our choice of PAC metric, we repeated our analyses using the modulation index described by Özkurt & Schnitzler (Özkurt-MI; 2011), which has been shown to provide similar results to the PLV-MI in the context of visually-induced PAC (Seymour et al., 2017, 2019). Results from the Özkurt-MI analysis were largely consistent with the PLV-MI analyses and are presented in Supplementary Materials.

We estimated PAC for every *f*_p_, *f*_a_ pair within two 700 ms time windows: a pre-stimulus window (-0.8 to -0.1 s relative to stimulus onset) and a post-stimulus window (0.3 to 1.0 s). These windows were selected to allow at least 200 ms of padding from the start and end of the epoch, to avoid edge artefacts induced by filtering (Kramer et al., 2008; Seymour et al., 2017). Our post-stimulus window also avoided the stimulus-induced evoked response. This procedure was implemented independently for each trial; estimates were then averaged across trials to produce one *f*_p_ × *f*_a_ grid of PAC estimates, known as a *comodulogram*, per time window (pre- and post-stimulus), and subject. Finally, PAC estimates were normalized using a surrogate analysis. Here, we estimated PAC for each *f*_p_, *f*_a_ pair after extracting phase and amplitude information from two randomly selected trials, respectively, and randomly shuffling the phase time series. This procedure served to destroy any true coupling by disrupting within-trial phase– amplitude correspondence and the temporal structure of the phase signal; the resulting PAC estimates therefore provided a baseline estimate of spurious PAC. We repeated this procedure 200 times, which generated 200 surrogate comodulograms. We then subtracted the mean surrogate comodulogram from the original pre- and post-stimulus comodulograms. All statistical analyses (described below) were performed on these surrogate-normalized comodulograms.

### 2.6. Gamma Power Computation

Since gamma power is highly correlated with age (Rhodes, Rier, Singh, et al., 2025), we estimated stimulus-elicited change in broadband (30-80 Hz) gamma power (hereafter simply “gamma change”) for each subject, to include as a covariate in our age-related analyses. Consistent with our PAC analysis, epochs were split into pre-stimulus (-0.8 to -0.1 s) and post-stimulus (0.3 to 1.0 s) windows, and data filtered to the centre frequency at each 2 Hz step of the 30-80 Hz range, using a two-pass fourth-order Butterworth filter (bandwidth = 4 Hz). We used a Hilbert transform to extract the absolute amplitude for each centre frequency (averaged across trials), for each time window. We calculated the normalized relative change between the post-stimulus and pre-stimulus windows ([post - pre] / pre); gamma change was the average relative change across frequencies.

### 2.7. Statistical Inference

Much of our statistical inference employed Bayes factors. Bayes factors provide an alternative to conventional null-hypothesis significance testing (NHST) and are commonly applied to group-level statistical inference on M/EEG data (e.g., Bailey et al., 2026; Bailey & Bardouille, 2025; Grootswagers, Robinson, & Carlson, 2019; Grootswagers, Robinson, Shatek, et al., 2019; Moerel et al., 2022, 2024; Proklova et al., 2019; Teichmann et al., 2022). A Bayes factor BF_10_ is a ratio of the conditional likelihood of some observed data under a model specifying an alternative hypothesis H_1_ (e.g., the true value of the parameter being tested is not equal to zero, θ ≠ 0) relative to a model specifying a null hypothesis H_0_ (e.g., θ = 0)^2^. BF_10_ values can therefore be interpreted as a quantitative measure of strength of evidence: BF_10_ < 1.0 indicates increasing support for H_0_ as values approach 0, and correspondingly BF_10_ > 1.0 indicates increasing support for H_1_ as values approach ∞ (Dienes, 2014, 2016; Schmalz et al., 2021). For reference, BF_10_ values exceeding 3.0 and 10.0 are conventionally labelled as “moderate” and “strong” evidence for H_1_ respectively (Dienes, 2014; Lee & Wagenmakers, 2014). When treated as a continuous measure of evidence, Bayes factors can be interpreted at face value and do not require correction for multiple comparisons (Dienes, 2016; Teichmann et al., 2022). Throughout this manuscript, we refer to Bayes factors quantifying evidence for multiple effects of interest: BF_Stimulus_ (quantifying evidence for a stimulus-related effect), BF_Age_ (for an age-related effect), and so on; in all cases, these notations refer to BF_10_ (that is, strength of evidence for a non-zero effect).

All Bayesian tests described in the following sections were implemented in the MATLAB environment using functions from the bayesFactor toolbox (Krekelberg, 2022) with default JZS priors (Liang et al., 2008; Rouder et al., 2012; Rouder & Morey, 2012).

#### 2.7.1. Detecting stimulus-induced PAC

To identify *f*_p_, *f*_a_ pairs exhibiting stimulus-induced increases in PAC (observed in Seymour et al., 2017, 2019; Voytek et al., 2010), we performed a paired Bayesian *t-*test (right tailed) at every cell of the comodulogram to compare PAC estimates from the pre-stimulus period to those of the post-stimulus period. Each test yielded a Bayes factor quantifying strength of evidence for a pre/post-stimulus difference: BF_Stimulus_. We implemented this procedure on data from children (*N* = 50) and adults (*N* = 51) separately; we reasoned that, if age-related changes in PAC were present, pooling data from both age groups may obscure potential PAC effects only present in one group. This analysis produced one *f*_p_ × *f*_a_ map of Bayes factors (henceforth BF maps) per age group. For completeness, we also conducted NHST nonparametric cluster-based testing, which yielded very similar results to our Bayesian clustering method (see Supplementary Materials).

#### 2.7.2. Detecting age-related changes in PAC

We investigated age-related changes in cells of the comodulogram exhibiting the strongest evidence for stimulus-induced PAC. To this end, we pooled BF_Stimulus_ values from both BF maps (children and adults) and identified the threshold for the top 5% of values (this was > 1.0 for both PAC metrics, confirming that this threshold enforced at least nominal support for H_1_). We then selected cells in each map exceeding the 5% threshold and identified clusters of contiguous suprathreshold cells. Finally, we created a binary mask from the largest cluster across both maps; this mask therefore represented the largest contiguous region of the comodulogram exhibiting the strongest evidence for stimulus-induced PAC, irrespective of age group. We note that this procedure yielded very similar results to non-parametric NHST-based clustering (see S Figure 3).

We subtracted subject-level pre-stimulus comodulograms from post-stimulus comodulograms, yielding a single matrix of PAC difference values (hereafter “PAC change”) per participant. For each subject, we then computed their mean PAC change within the BF-derived logical mask. These values therefore represented the pre-/post-stimulus PAC difference at frequency pairs where stimulus-induced PAC was most evident at the group level.

We quantified the effects of age and gamma change on PAC change via covariate testing on Bayesian linear regression models (Rouder & Morey, 2012). Within a Bayesian multiple regression framework, Bayes factors for individual predictors can be obtained through model comparison: a model which includes the variable of interest (e.g., age) is compared to a model which does not include that variable. The ratio of marginal likelihoods under the two models (with the full model in the numerator) therefore gives the Bayes factor for the effect of that variable. Equivalently, one may compute a Bayes factor for each model (relative to an intercept-only model) and, in turn, the ratio between them (Rouder & Morey, 2012). Using this approach, we obtained Bayes factors for the linear effects of age and gamma change: respectively, BF_Age_ and BF_Gamma_.

As our age distribution was bimodal and Bayesian regression on ranked data would require deviation from default priors, we conducted, as a robustness check, a non-Bayesian (NHST) linear regression on ranked values, with ranked PAC change as the dependent variable and ranked age and gamma change as predictors.

## 3. Results

### 3.1. Stimulus-Induced PAC

We first investigated changes in PAC across pre- and post-stimulus time windows. We conducted a Bayes t-test at each cell of a comodulogram (i.e., grid of *f*_p_, *f*_a_ pairs) to compare PAC between the two time windows. This produced a *f*_p_ × *f*_a_ grid with a Bayes factor (BF_Stimulus_) at each cell; higher values indicate stronger evidence for increased PAC in the post- (relative to pre-) stimulus window. We performed these comparisons independently for children and adults.

Figure 2 shows grand-average PAC change and Bayes factors. For the child group, there was a numerical effect (visible in Figure 2A) whereby gamma amplitude between 50-80 Hz was modulated by low-frequency phase between 10-12 Hz. However, statistical evidence for this effect was relatively weak, with maximum BF_Stimulus_ = 2.72 (within the range of weak/anecdotal evidence; Lee & Wagenmakers, 2014. Moreover, parallel NHST analyses revealed no significant clusters in the child group: see S Figure 3). By contrast, in the adult group, there was a clear stimulus-induced increase in PAC whereby gamma amplitude around 50-70 Hz was modulated by low-frequency phase between 6-10 Hz.

**Figure 1.**
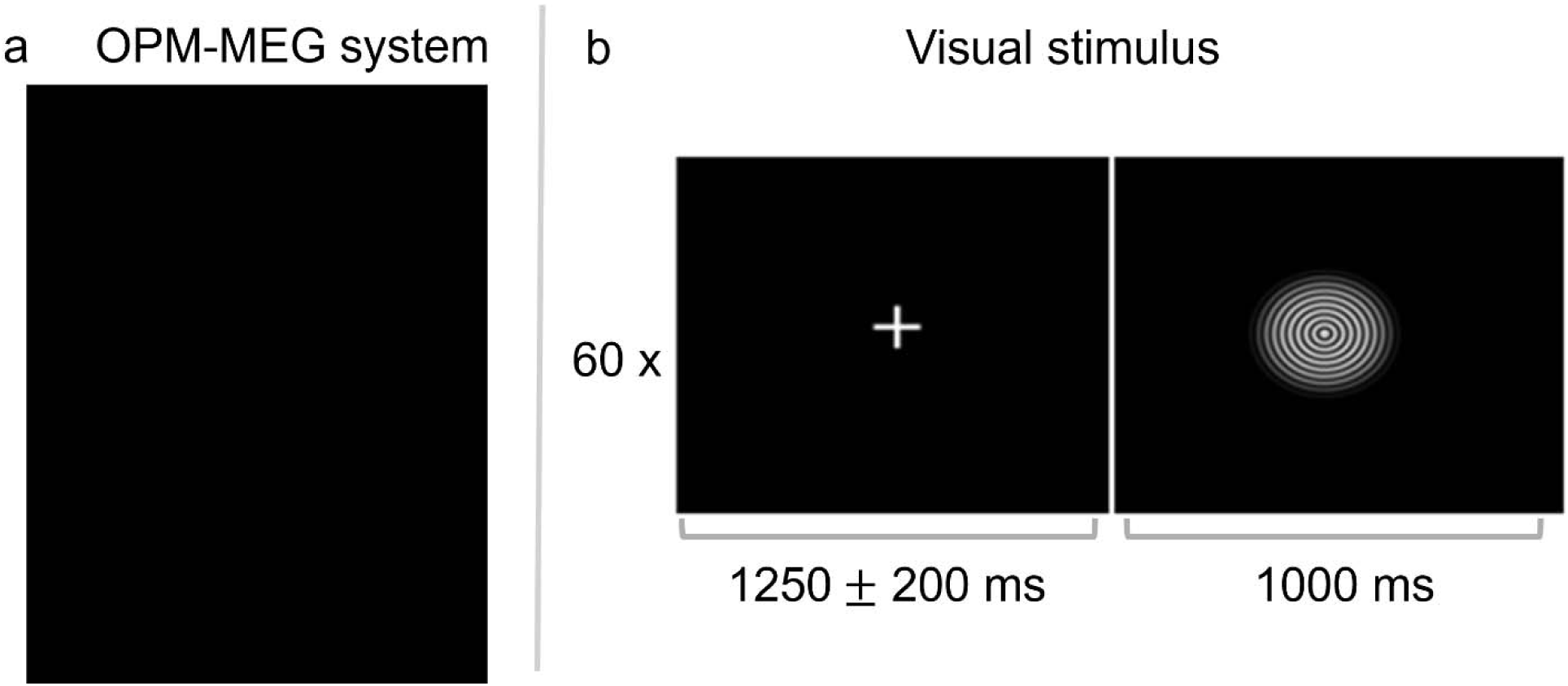
Left panel (a): a child in the OPM-MEG system used in this study, at the SK site. **This panel has been occluded to comply with bioRxiv policy.** Right panel (b): Frames from a single trial in the visual stimulation paradigm. Figure reproduced from Rhodes, Rier, and Singh, et al., (2025).

**Figure 2.**
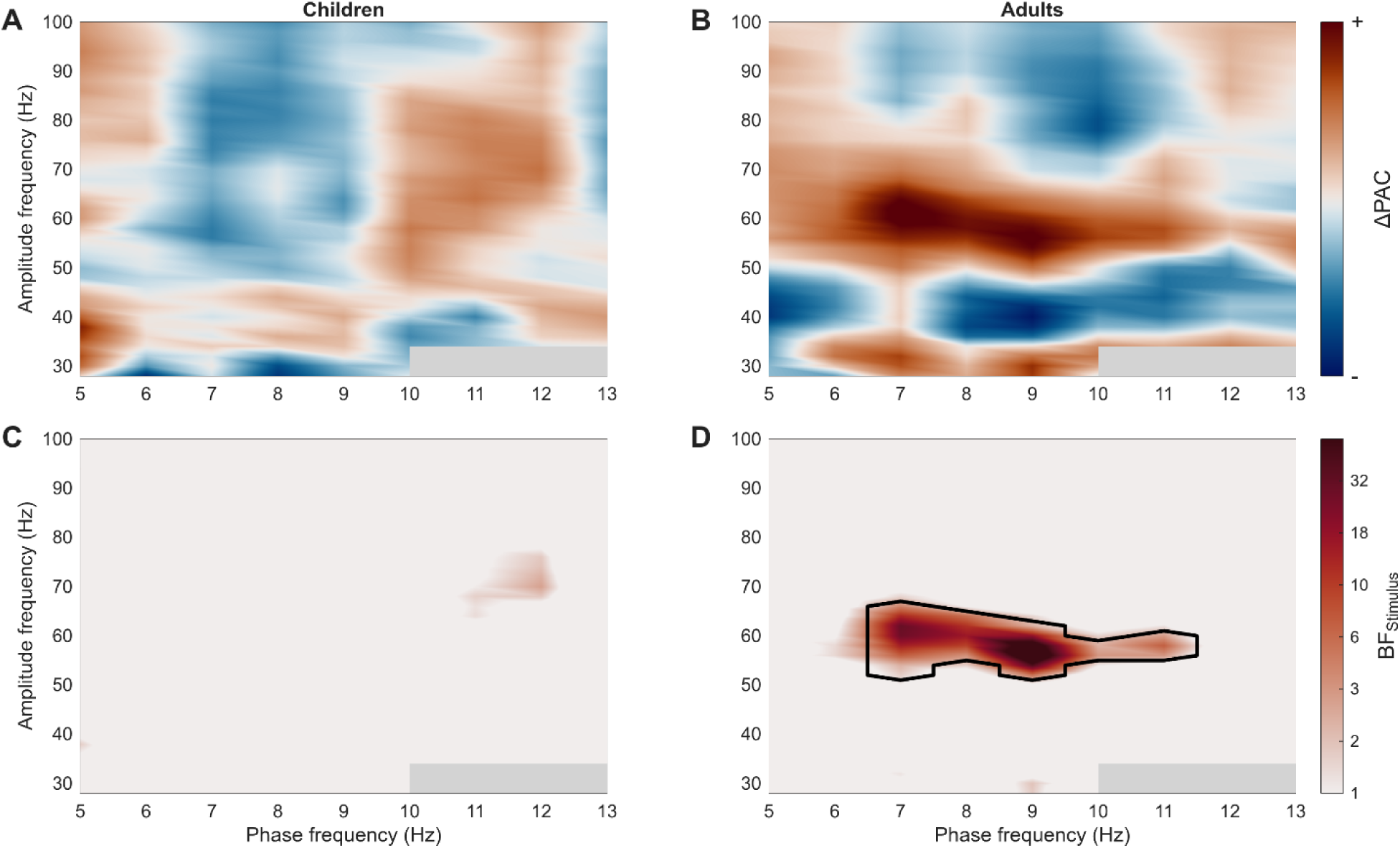
Stimulus-induced PAC separately for children (**A, C**) and adults (**B, D**). Top row shows grand-average comodulograms for the post-stimulus minus pre-stimulus difference. Botton row shows Bayes factors (BF_Stimulus_) comparing PAC in the post-stimulus window to that of the pre-stimulus window. Plotted BF values are log10- transformed for scaling; the colour bar shows back-transformed (i.e., real BF_Stimulus_) values for interpretability. The black contour indicates the largest contiguous cluster of cells in the top 5% of BF_Stimulus_ values across both age groups. Values at the intersection of 10-13 Hz and 28-34 Hz are masked out (grey rectangles) due to our insensitivity to PAC in this range.

**Figure 3.**
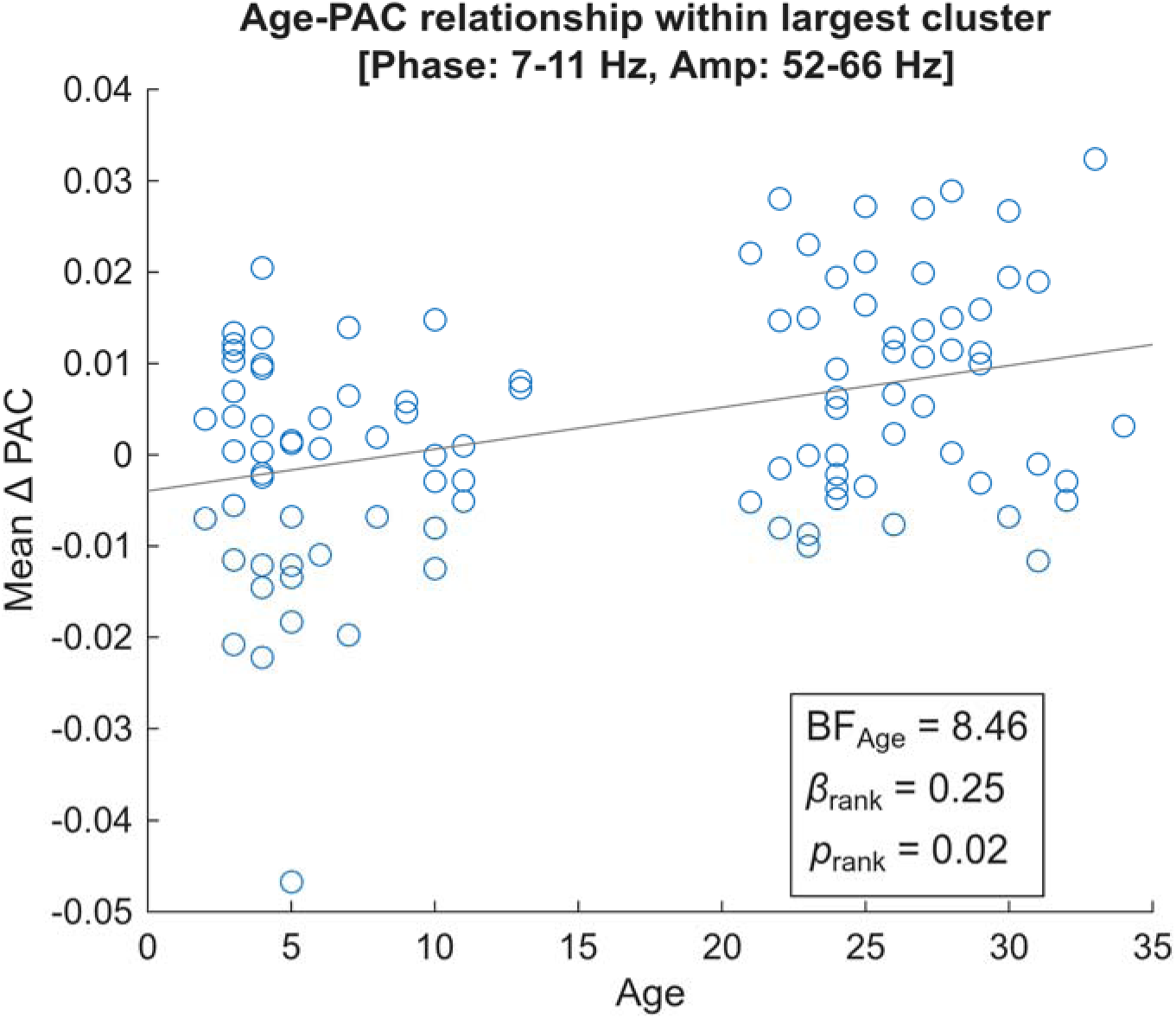
Scatterplot shows the relationship between age and the magnitude of stimulus-induced PAC change (post-stimulus minus pre-stimulus values), averaged within the group-level PAC cluster (see Figure 2D). BF_Age_ is the Bayes factor for the linear age effect with unranked data; β*_rank_* and *p*_rank_ were obtained from non-Bayesian linear regression on ranked values.

We identified the largest contiguous cluster of cells within the top 5% of BF_Stimulus_ values across both age groups (see Methods). This revealed a cluster in the adult comodulogram spanning a low frequency range of 7-11 Hz and a high frequency range of 52-66 Hz. This cluster was converted to a binary mask and carried through to the age-related analyses.

### 3.2. Age- and Gama-Related Effects

For each participant, we computed mean PAC change across cells within the high-evidence mask described above. We assessed complementary effects of age and stimulus-induced broadband gamma change on PAC change.

Covariate testing on Bayesian linear regression models (including age and gamma change as predictors) revealed moderate-to-strong evidence for an effect of age on PAC change (BF_Age_ = 8.46). This result was verified by a non-Bayesian linear regression on ranked values, which revealed a significant positive relationship between age and PAC change: β_rank_= 0.25, *p*_rank_ = 0.02. This age-PAC relationship is shown in Figure 3. By contrast, gamma change did not have a clear effect on PAC: BF_Gamma_ = 0.34, β_rank_ = 0.11, *p*_rank_ = 0.27.

## 4. Discussion

We investigated developmental changes in PAC elicited by a visual stimulation paradigm, in a cohort of children and adults aged 2-34 years. Consistent with previous work (Seymour et al., 2017, 2019; Voytek et al., 2010), the adult group exhibited stimulus-related coupling between the phase of oscillations in the high theta / alpha range (6-11 Hz) and the amplitude of oscillations in the gamma range (52-66 Hz). Meanwhile, PAC was absent in the child group. We observed a positive relationship between age and visual PAC which, importantly, was robust when controlling for gamma power. We also note that the visually-induced gamma ERS response was detectable in this group of children (Rhodes, Rier, Singh, et al., 2025). These findings are consistent with previous work showing population-level changes in PAC despite no differences in within-band oscillatory power (Khan et al., 2013; Seymour et al., 2019), and suggest that gamma ERS and visual PAC may represent developmental dissociable processes, with distinct maturational trajectories.

In the context of visual processing, oscillations in the gamma band are thought to support bottom-up processing of sensory input (e.g., Bastos et al., 2015), while PAC has been linked to top-down attentional control of this process (Trajkovic et al., 2025). In turn, this top-down control is likely relayed to visual cortex via long-range, low-frequency phase synchrony with prefrontal regions (Trajkovic et al., 2025) and possibly occipital regions further up the visual processing hierarchy (Seymour et al., 2019). Our findings may reflect gradual maturation of top-down control over perceptual processing, as opposed to relatively rapid development of bottom-up processing (evident from the presence of gamma ERS in young children). An interesting avenue for further work may be to explore the relation between long-range alpha connectivity and visual PAC, and whether this strengthens with age.

We were, however, surprised that we did not detect PAC in the child group. To our knowledge, two studies have reported stimulus-related PAC effects in school-aged children (> 7 years of age; Barnes et al., 2016; Mamashli et al., 2021); by contrast, our sample of children was considerably younger (over half were < 6 years old), and so our results are not directly comparable to those studies. Interestingly, Attaheri et al. (2022) detected delta-gamma and theta-gamma PAC in infants (ages 4-11 months) listening to nursery rhymes, using whole-head EEG. However, these authors report that the magnitude of PAC did not vary with age, while the phase-carrying frequencies appeared to reflect neural sensitivity to properties of the auditory stimulus (i.e., they were elicited exogenously), rather than arising endogenously to support top-down perceptual control. These considerations highlight the fact that PAC is a versatile form of neural signaling whose functional role depends largely on the context in which it is observed. As such, PAC observed in one experimental setting is not necessarily comparable to that of other settings.

Our study opens the door to a few interesting lines of investigation for future developmental research. A lingering question is whether age-related changes in visual PAC are strictly linear, or whether PAC is subject to rapid development over some critical period (e.g., during adolescence, considering that visual PAC has been observed in a group of mid-to-late adolescents and young adults; Seymour et al., 2019). Further research might investigate potential non-linear PAC changes over a uniform sample of ages (comprising children, adolescents, and adults). Moreover, some work has demonstrated that PAC is disrupted in neurodevelopmental disorders such as autism (e.g., Khan et al., 2013; Mamashli et al., 2018; Peck et al., 2022; Seymour et al., 2019). It is possible that this population also exhibits altered developmental trajectories of visual PAC compared to neurotypical controls. Research in this vein may further our understanding of (and ability to detect) early neurodevelopmental markers of autism.

In summary, our results support a nuanced view of functional development of the visual system. As highlighted in previous work (Bastos et al., 2015; Trajkovic et al., 2025), visual perception is not a simple, passive process, but entails multiple mechanisms (both bottom-up processing and top-down control), which in turn are organized across multiple frequency bands and spatial scales. Our findings extend those of previous work reporting age-related changes in within-band responses to visual stimuli (Gaetz et al., 2011; Rhodes, Rier, Singh, et al., 2025); here we showed that visual PAC is similarly age-dependent, but may emerge relatively late during development. Subsequent work might further interrogate the (non)linearity of this effect, or its viability as a marker for neurodevelopmental disorders.

## Data and code availability statement

Data from UoN is available on Zenodo (https://zenodo.org/records/15190020), data from SK will be made available through Ontario Brain Institute. Code for the analyses presented in this manuscript is available on GitHub: https://github.com/lyambailey/Visual_PAC_OPM_2026.

## Abbreviations

BF: Bayes factor
ERS: Event-related synchronization
MEG: Magnetoencephalography
OPM: Optically-pumped magnetometer
PAC: Phase-amplitude coupling
SK: SickKids (data collection site)
UoN: University of Nottingham (data collection site)

## Acknowledgements

We would like to thank all our participants and the families of our young participants for their involvement in this study.

## CRediT contributions

L.M.B. Conceptualization, Visualization, Writing - original draft, Writing - review & editing, Formal analysis, Methodology, Software, Data curation. N.R. Investigation, Writing - review & editing, Conceptualization, Data curation, Project administration. R.A.S. Writing - review & editing, Conceptualization. M.J.B. Funding acquisition, Supervision, Writing - review & editing, Project administration, Resources. M.J.T. Supervision, Funding acquisition, Writing - review & editing, Project administration, Resources, Conceptualization.

## Footnotes

1 This procedure led to numerical instability at lower phase frequencies. We therefore implemented Fieldtrip’s cfg.bpinstabilityfix = ’reduce’ option which, in cases of instability, automatically reduced the filter order until a stable filter could be constructed.

2 Note that the "10" in BF_10_ denotes that likelihood under the alternative hypothesis (H_1_) is in the numerator of the ratio and likelihood under the null (H_0_) is in the denominator; therefore values > 1.0 favour H_1_. Bayes factors can alternatively be represented as BF_01_ (whereby the ratio is flipped), in which case values > 1 indicate support for the null.

