## Supplementary Materials for "OPM-MEG reveals protracted maturation of visual phase-amplitude coupling from early childhood to adulthood"

**Re-Analysis with Özkurt-MI**

Our main analyses quantified PAC using the phase-locking-value modulation index (PLV-MI). To verify that our results were robust to the choice of PAC metric, we repeated our analyses using the modulation index described by Özkurt & Schnitzler (2011), hereafter Özkurt-MI. The Özkurt-MI assumes that, if PAC is present, high amplitude values (computed from *f*_a_) will occur preferentially at particular phases (from *f*_p_), as opposed to being uniformly distributed across the phase cycle. Here, continuous data are represented by a time-varying complex-valued signal comprising an angle component (i.e., phase information) and an amplitude component. In this case, the angle and amplitude components were computed from *f*_p_ and *f*_a_ respectively via Hilbert transform. Each time point is represented by a single vector in this complex-valued space. The mean vector length over timepoints provides a "raw" modulation index, characterizing the extent to which amplitude is distributed across possible phases (e.g., a mean vector length of zero indicates that amplitude is uniformly distributed across the phase cycle) (Canolty et al., 2006). This measure of PAC can be affected by *f*_a_ power, therefore the Özkurt-MI additionally normalizes the mean vector length according to the power of *f*_a_ (Özkurt & Schnitzler, 2011).

All other analysis steps (comparison of pre-stimulus to post-stimulus comodulograms, Bayes factor computation and clustering, and analyses of age- and gamma-related effects) were identical to those described in the manuscript proper.

**Stimulus-Induced PAC**

S Figure 1 shows grand-average PAC difference values (PAC change: post-stimulus minus pre-stimulus) and Bayes factors (BF_Stimulus_) for the child and adult group. As in the main (PLV-MI) analyses, there was only weak evidence for stimulus-induced PAC within the child group (Max BF_Stimulus_ = 1.59). In the adult group, stimulus-induced PAC was evident between phase frequencies of 6-9 Hz and amplitude frequencies of 50-70 Hz. The largest cluster of values in the top 5% across both age groups revealed a cluster in the adult group, spanning phase frequencies 6-9 Hz and amplitude frequencies 50-68 Hz.

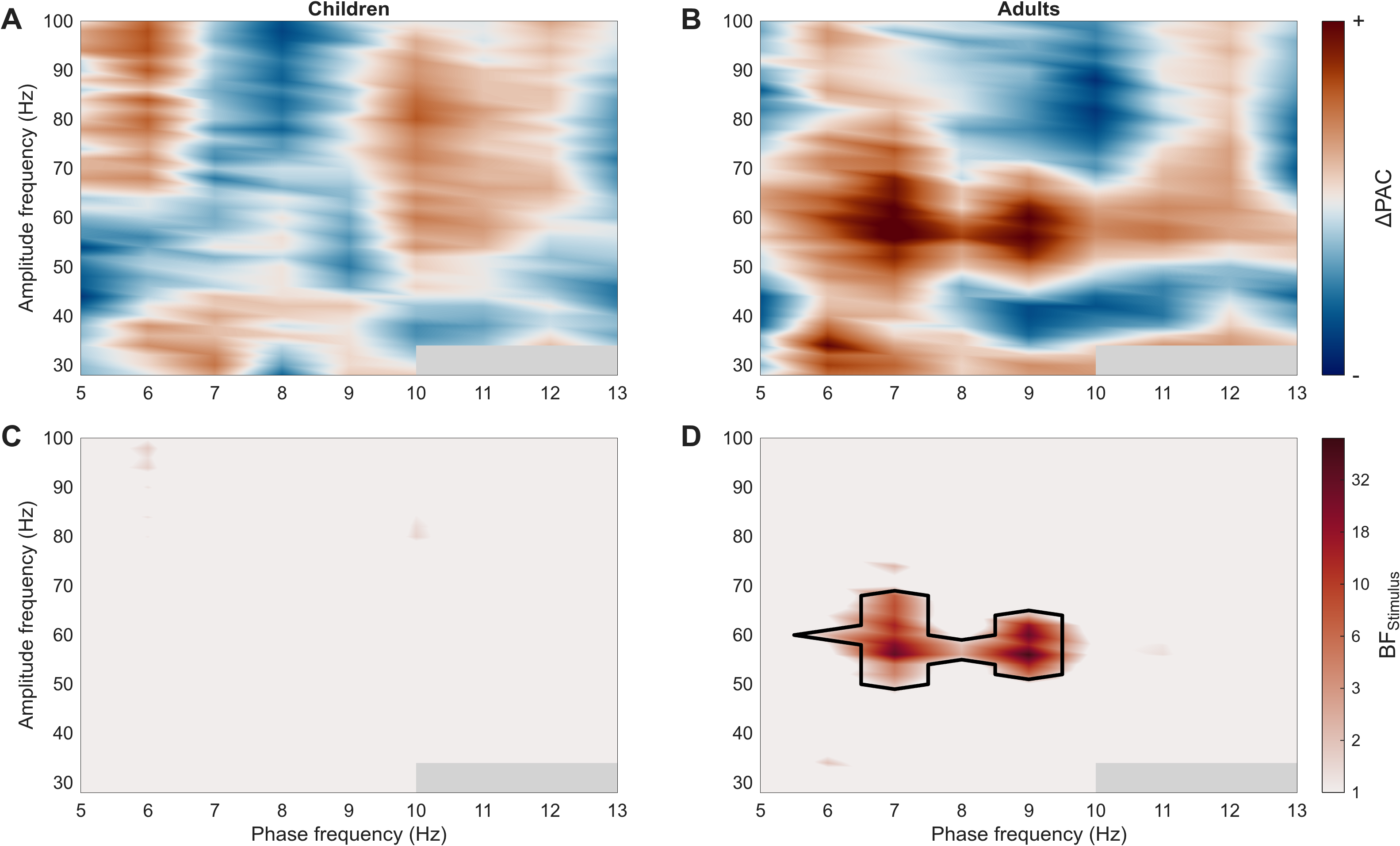

*S Figure 1*. Stimulus-induced PAC (as measured by the Özkurt-MI) separately for children (**A, C**) and adults (**B, D**). Top row shows grand-average comodulograms for the post-stimulus minus pre-stimulus difference. Botton row shows Bayes factors (BF_Stimulus_) comparing PAC in the post-stimulus window to that of the pre-stimulus window. Plotted BF_Stimulus_ values are log10-transformed for scaling; the colour bar shows back-transformed (i.e., real BF_Stimulus_) values for interpretability. The black contour indicates the largest contiguous cluster of cells in the top 5% of BF_Stimulus_ values across both age groups. Values at the intersection of 10-13 Hz and 28-34 Hz are masked out (grey rectangles) due to our insensitivity to PAC in this range.

**Age- and Gamma-Related PAC**

Subject-level PAC change values were averaged across cells within the high-evidence cluster shown in S Figure 1D. We assessed complementary effects of age and stimulus-induced broadband gamma change on PAC change. Bayesian covariate testing on unranked values revealed anecdotal evidence for an age effect on PAC change, BF_Age_ = 2.44. While the support for this effect was weaker than the corresponding PLV-MI age effect (BF_Age_ = 8.46), it replicated the direction and, also consistent with PLV-MI, was statistically significant following non-Bayesian linear regression on ranked values, *β* = 0.22, *p* = 0.04. This effect is shown in S Figure 2. Meanwhile, there was no clear effect of gamma change on PAC change, BF_Gamma_ = 0.61, *β* = 0.21, *p* = 0.051.

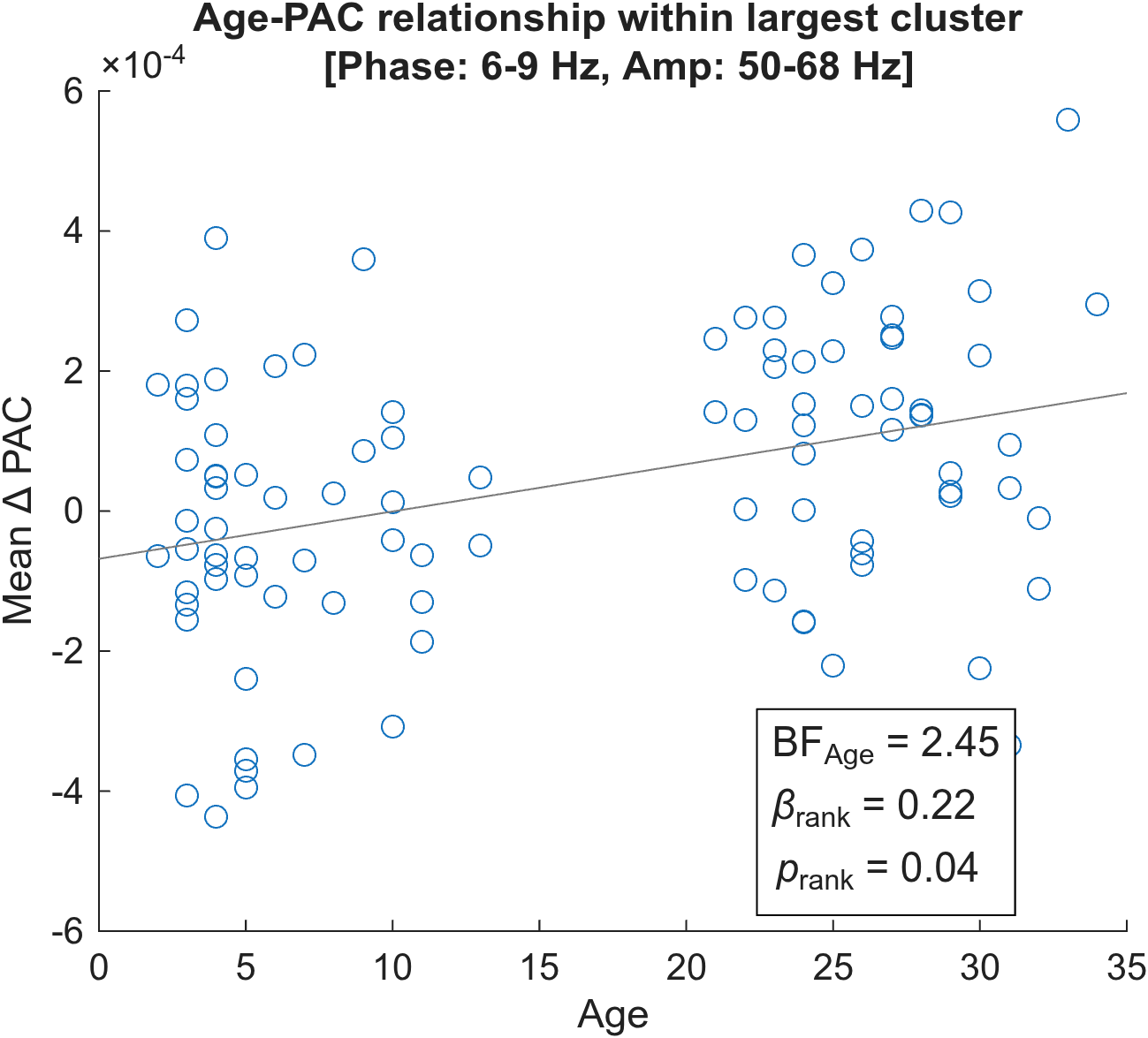

*S Figure 2*. Scatterplot shows the relationship between age and the magnitude of stimulus-induced PAC change, as measured by the Özkurt-MI (post-stimulus minus pre-stimulus values), averaged within the group-level PAC cluster (see S Figure 1D). BF_Age_ is the Bayes factor for the linear age effect with unranked data; *β_rank_* and *p*_rank_ were obtained from non-Bayesian linear regression on ranked values.

Overall, results from the Özkurt-MI analysis were consistent with those of PLV-MI in terms of the frequency ranges at which stimulus-induced PAC occurred (phase frequencies ~6-10 Hz, amplitude frequencies ~50-70 Hz), the significant positive relationship between age and PAC change, and the absence of gamma-related effects.

**Detecting stimulus-induced PAC with NHST**

The following procedures were performed for the PLV-MI and Özkurt-MI measures. Each participant was associated with two comodulograms for the pre- and post-stimulus periods, respectively. We used non-parametric cluster-based testing to detect stimulus-induced increases in PAC (separately in children and adults, as in our main analyses) using MATLAB functions adapted from publicly available code (https:// github.com/neurofractal/sensory_PAC). At each cell of the comodulogram we conducted a right-sided paired-sampled *t*-test to compare post-stimulus values to pre-stimulus values. This produced a *t* value at each cell, and clusters were formed from contiguous cells whose *t* values were significant at the *p* < 0.05 level (uncorrected). Cluster-level *t* values (maximum within each cluster) were compared to a null distribution that was constructed by randomizing condition labels (pre-/post-stimulus) and repeating t-tests (1,000 permutations); the null distribution comprised maximum cluster-level *t*-values from each permutation. Original clusters whose *t* value exceeded the top 5% of the null distribution were deemed significant.

Both measures revealed a significant cluster in the adult group (see S Figure 2); these clusters were of similar spatial extent to those detected by our Bayesian analyses. No significant clusters were found in the child group, for either measure.

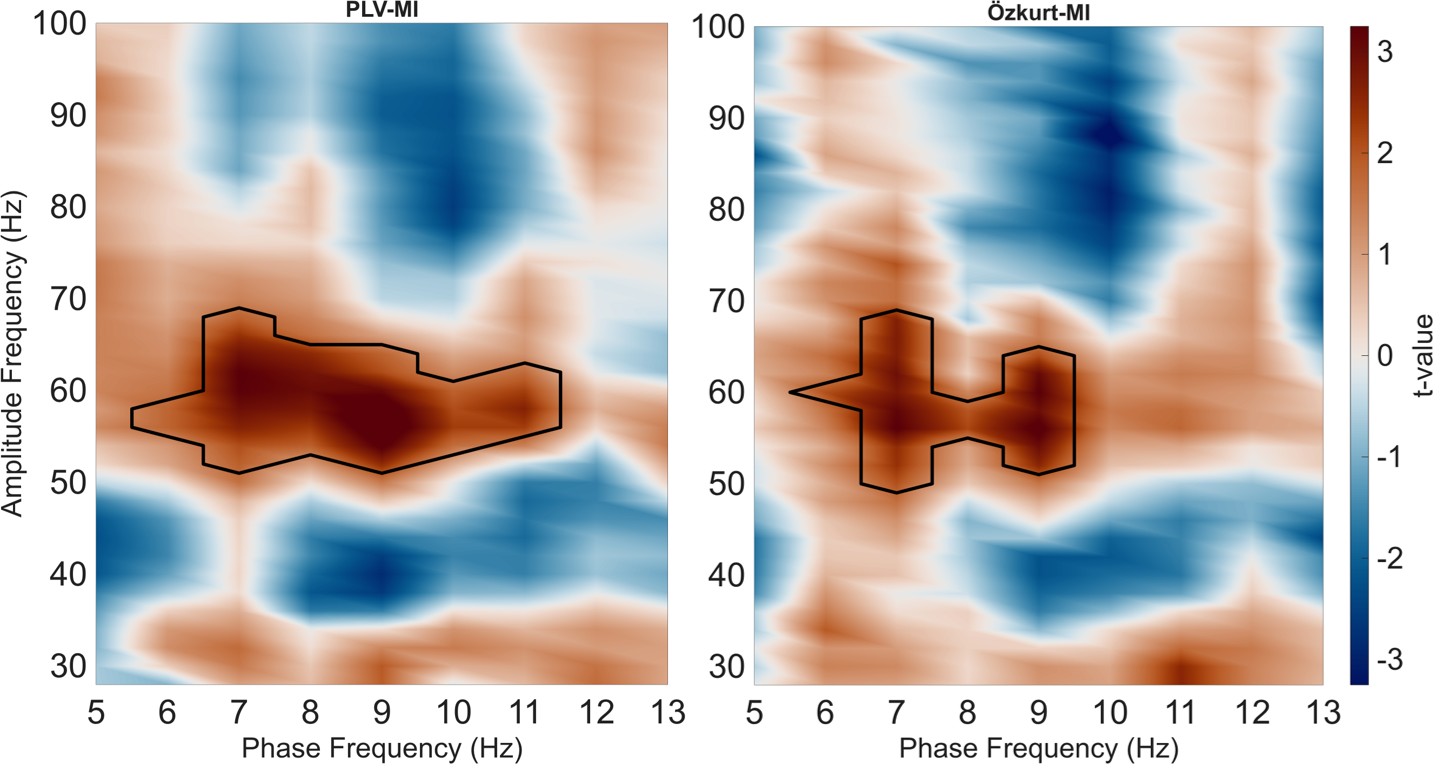

*S Figure 3.* Results of NHST nonparametric testing, comparing PAC in the post-stimulus period to that of the pre-stimulus period, in the adult group only (no significant clusters were present in the child group). Results are shown for the PLV-MI (left panel) and Özkurt-MI (right panel) measures. Black contours show significant clusters (*p* < 0.05) revealed by non-parametric cluster-based statistics.
